# Population receptive fields of human primary visual cortex organised as DC-balanced bandpass filters

**DOI:** 10.1101/593608

**Authors:** Daniel Gramm Kristensen, Kristian Sandberg

**Author notes:** **Corresponding author**, Daniel Gramm Kristensen, Center of Functionally Integrative Neuroscience, Aarhus University Hospital, Nørrebrogade 44, Building 1A, 8000 Aarhus C, Denmark.

## Abstract

The response to visual stimulation of population receptive fields (pRF) in the human visual cortex can be accurately modelled with a Difference of Gaussians model, yet many aspects of their organisation remain poorly understood. Here, we examined the theoretical underpinnings of this model and argue that the DC-balanced Difference of Gaussians (DoG) holds a number of advantages over a DC-biased DoG. Through functional magnetic resonance imaging (fMRI) pRF mapping, we compared performance of DC-balanced and DC-biased models in human primary visual cortex and found that when model complexity is taken into account, the DC-balanced model is preferred. Finally, we present evidence indicating that the BOLD signal DC-offset contains information related to the processing of visual stimuli. Taken together, the results indicate that V1 neurons are at least frequently organised in the exact constellation that allows them to function as bandpass-filters, which allows for the separation of stimulus contrast and luminance. We further speculate that if the DoG models stimulus contrast, the DC-offset may reflect stimulus luminance. These findings suggest that it may be possible to separate contrast and luminance processing in fMRI experiments and this could lead to new insights on the haemodynamic response.

## Introduction

Neurons in the early visual system have a central, excitatory receptive field as well as a surrounding, inhibitory receptive field. Visual stimuli presented in the central receptive field cause the neuron to respond by increased firing whereas stimuli presented in the surround are by themselves subthreshold but can modulate the response to a central stimulus^1–4^. The dynamics of the centre-surround constellation can be examined using functional magnetic resonance imaging (fMRI), and it has been demonstrated that a Difference of Gaussians (DoG) population Receptive Field (pRF) model yields improved predictions compared to a single Gaussian pRF model^5^. In this experiment, we used the following mathematical description of the DoG model:

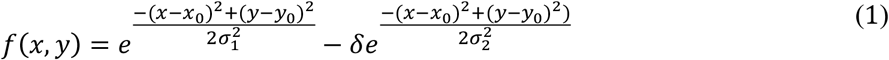

where x and y are the Cartesian coordinates of the visual field with x_0_ and y_0_ designating the location of the receptive field centre. σ_1_ and σ_2_ are the spatial extent of the two Gaussians and δ is a parameter that modulates the amplitude of the second inhibitory Gaussian. The DoG is sometimes characterised as an edge-detector, but more precisely, it can be said that the DoG detects *changes* of certain frequencies, i.e. the DoG detects changes in contrast. In natural scenes, these changes in contrast occur over a vast range of scales, which makes it difficult to find a single filter that is optimal at each scale. As a solution, it was therefore proposed to measure luminance and contrast separately at various scales^6^, but the neural implementation of this remains unclear. Luminance is here defined as the local average intensity, meaning the average over the receptive fields’ spatial extent. The average of a signal is also referred to as its DC-offset, and so for the DoG to strictly measure changes in contrast it must be DC-balanced. To establish whether a given DoG is DC-balanced or not, we make the following observations. We reason the DoG is DC-balanced when the double integral of the 2D spatial filter equals zero, such that

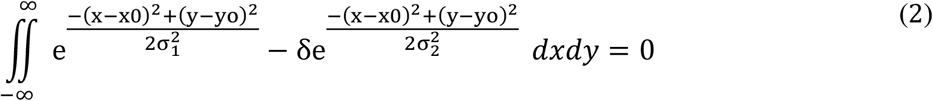

We assume *σ*_2_ > 0 and *σ*_1_ > 0 and that the Gaussians are circularly symmetric and centered on the same 2D Cartesian coordinate. The integral of a 1D Gaussian function is then given by

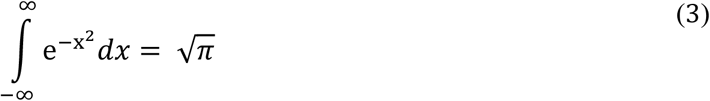

so that

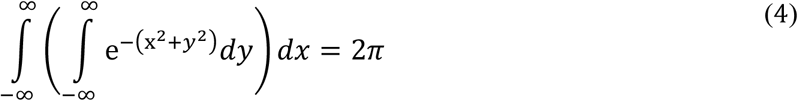

which for equation (2) yields

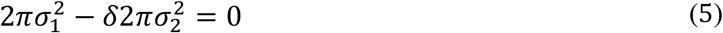

Thus

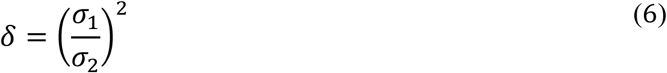

In the following, we therefore distinguish between a DC-balanced DoG and a DC-biased DoG. A DC-biased DoG can be either positively biased or negatively biased, but not both. The DoG model is positively biased when its total energy is greater than zero and negatively biased when it is lower than zero. Equation 6 also reveals that when DC-balance is enforced and δ approaches 1, σ_1_ and σ_2_ will have to compensate by approaching the same value. This is easily demonstrated by rewriting equation 6 to solve for σ_1_ instead of δ, which gives 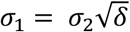. As δ approaches 1, 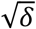 also approaches 1 and so σ_1_ approaches σ_2_. However, as the size of the two Gaussians approaches one another, the amplitude of the central Gaussian of the DoG approaches zero. On the other hand, if we solve for σ_1_, and let δ approach zero, σ_2_ approaches infinity. Finally, if *δ* = (*σ*_1_/*σ*_2_)^2^ and δ > 1, then it follows that *σ*_1_ > *σ*_2_. In these cases, the DoG has been turned upside down so it instead consists of an inhibitory central Gaussian and a surrounding excitatory Gaussian. In the following, we have therefore restricted our model space to 0.1 < δ < 0.9.

The changes the DoG is capable of detecting depends on the specific values of δ, σ_1_ and σ_2_. To demonstrate these properties, we have in Figure 1A plotted three sinusoidal signals with a frequency of 0.1 cycles per unit distance (c/d), 0.03 c/d and 0.05 c/d in arbitrary units. The signal with a frequency of 0.05 c/d has a DC-offset of 0, meaning no DC-offset. The signal with a frequency of 0.03 c/d has a DC-offset of 3.7 and in addition a low frequency oscillation. The signal with a frequency of 0.1 c/d has a DC-offset of 9.2 and also exhibits a linear trend as a function of distance. Figure 1B shows an alternative definition of the DC-offsets calculated using a running average with a window size of 100. This is included to show that the definition of the DC-offset is contingent on the window over which it is analysed. Figure 1C and 1D shows the result of convolving the three signals with a positively DC-biased DoG and a DC-balanced DoG, respectively. The two models are shown in the inset of Figure 1D in both the spatial domain (left) and spatial frequency domain (right). Here, the DC-biased model is plotted in grey, whereas the DC-balanced model is plotted in black.

**Figure 1.**
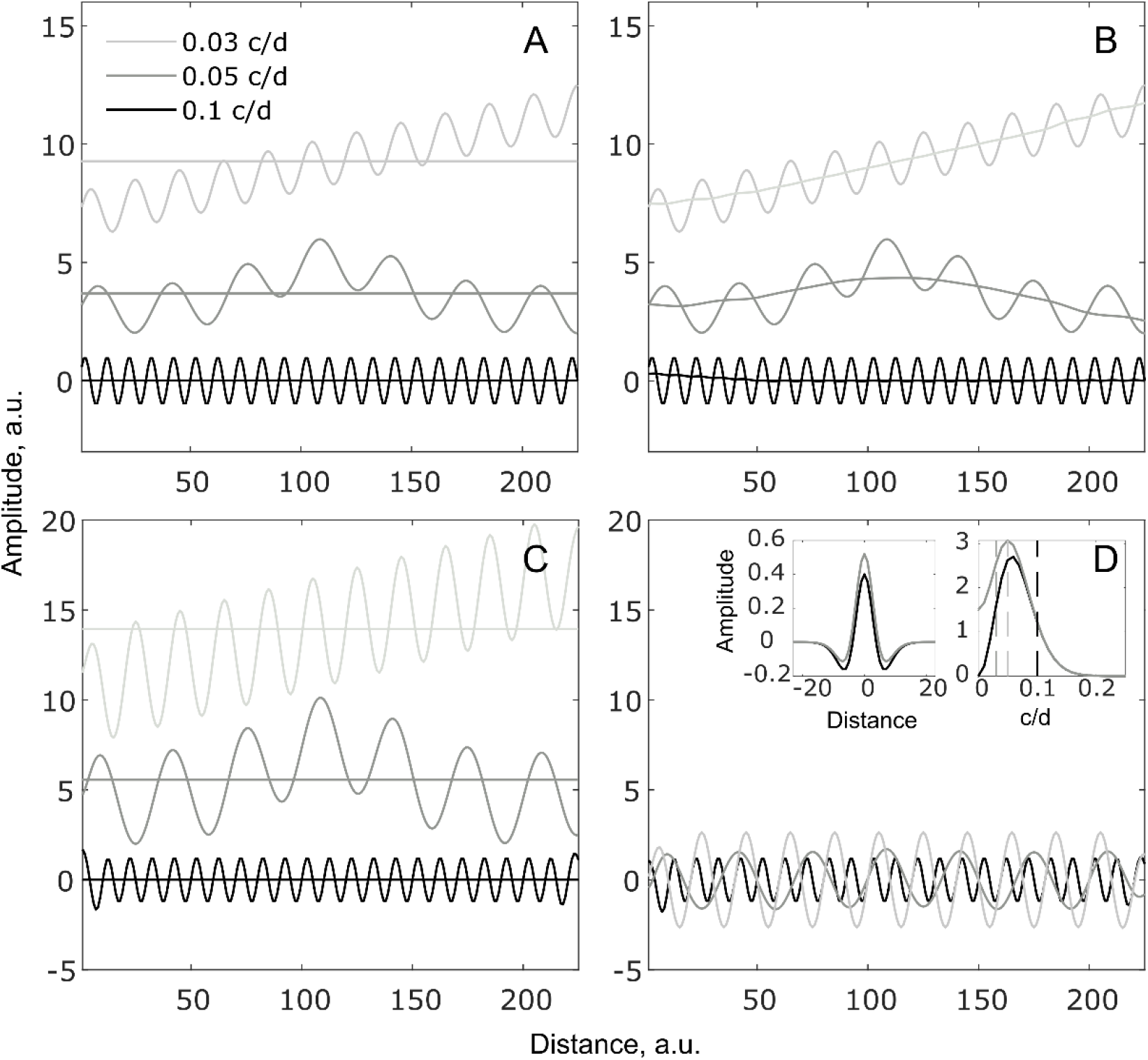
Effects of filtering sinusoidal signals with Difference of Gaussians. **A:** Three sinusoidal signals with frequencies as shown in the plots. All signals have the same amplitude. Straight lines indicate their DC-offset. **B:** Running average of window size 100 calculated for the 3 signals. **C:** Sinusoidal signals after convolution with a positively DC-biased DoG. **D:** Same as C, but for DC-balanced DoG. **Inset, left:** DoG models in the spatial domain. **Inset, right:** DoG models in the frequency domain. The dashed lines indicate the frequency of the sinusoidal signals.

Note that the scale on the y-axis is different between the two upper figures and the two lower figures. Figure 1C illustrates that for the two DC-offset signals, the DC-biased DoG model retains the DC-offset, the signal drift, and the low frequency oscillation, whereas Figure 1D shows this is not the case for the DC-balanced DoG. Common to both models is that the signals’ amplitudes change after convolution. The cause of this is shown in the rightmost inset of Figure 1D, where the two models’ frequency sensitivity is shown. Here, it can be seen that a given DoG model is not equally sensitive to all frequencies and that the DC-biased model is a lowpass filter, whereas the DC-balanced model is a bandpass filter. Furthermore, the two models have the same sensitivity to frequencies of 0.1 c/d, whereas the DC-biased model is more sensitive to both frequencies of 0.03 c/d as well as 0.05 c/d. We have refrained from normalising the amplitude of the frequency response to better illustrate this. Finally, in Figure 1C it is also seen that the DC-offset of the two signals that had a DC-offset has increased to about 5.5 and 14 because of convolution with the DC-biased model, whereas the signal with no DC-offset remains without a DC-offset. The DC-offset increase is due to the DC-biased model being positively biased and illustrates that the DC-biased model cannot separate luminance and contrast.

There is evidence that the excitatory and inhibitory parts of single cell receptive fields tend to balance out^7^. This suggests that single-cell receptive fields are DC-balanced. It has also been shown that luminance is largely independent of contrast in natural images^8^. This is in line with the proposition that contrast and luminance may be processed by separate mechanisms^6^. We therefore investigated whether population receptive fields of human V1 could be said to be DC-balanced by conducting a population Receptive Field mapping experiment^9^ with 22 healthy human participants. We chose to focus on V1 because it yielded a higher spatial resolution compared to mapping the entire visual cortex. If pRFs of V1 are indeed DC-balanced, it suggests that the variance explained by the DoG model is primarily concerned with stimulus contrast. We therefore also conducted a preliminary analysis to see if it was possible to identify information in the BOLD signal related to stimulus luminance.

## Results

Participants of the experiment were placed in the bore of a Siemens Magnetom 3T Skyra. Through a mirror, they were presented with expanding rings simultaneous with clockwise rotating wedges or contracting rings simultaneous with counter-clockwise rotating wedges, both with high-contrast ripple-like textures^10^. Example frames of this stimulus texture are supplied in Supplementary Figure S1 and S2. We also acquired high-resolution T1 structural images for each participant and used these for an anatomical reconstruction in Freesurfer and realignment of functional images. A delineation of each participant’s primary visual cortex (V1) was made from a prediction using landmarks from the anatomical reconstruction using a built-in Freesurfer algorithm^11^. We chose to use the anatomical predictions as they are less prone to human bias and they showed good agreement with functional delineation of V1, as has also been found previously^12^.

We then fitted two models to the obtained data; a DoG model where the δ-parameter was allowed to take any value and a DoG model where the δ-parameter was fixed in accordance with equation 6, and we therefore refer to the two models as the unrestricted model and the DC-balanced model. Other than this difference, the two procedures were identical and progressed as follows. First, we found the best correlation between a coarse parameter space and the observed data. This coarse fit served as the initial model parameters for the fine fitting procedure as illustrated in Figure 2. For each voxel, a binary representation of the stimulus at a given repetition time (TR) was multiplied with the initial DoG model parameters and the result summed.

**Figure 2.**
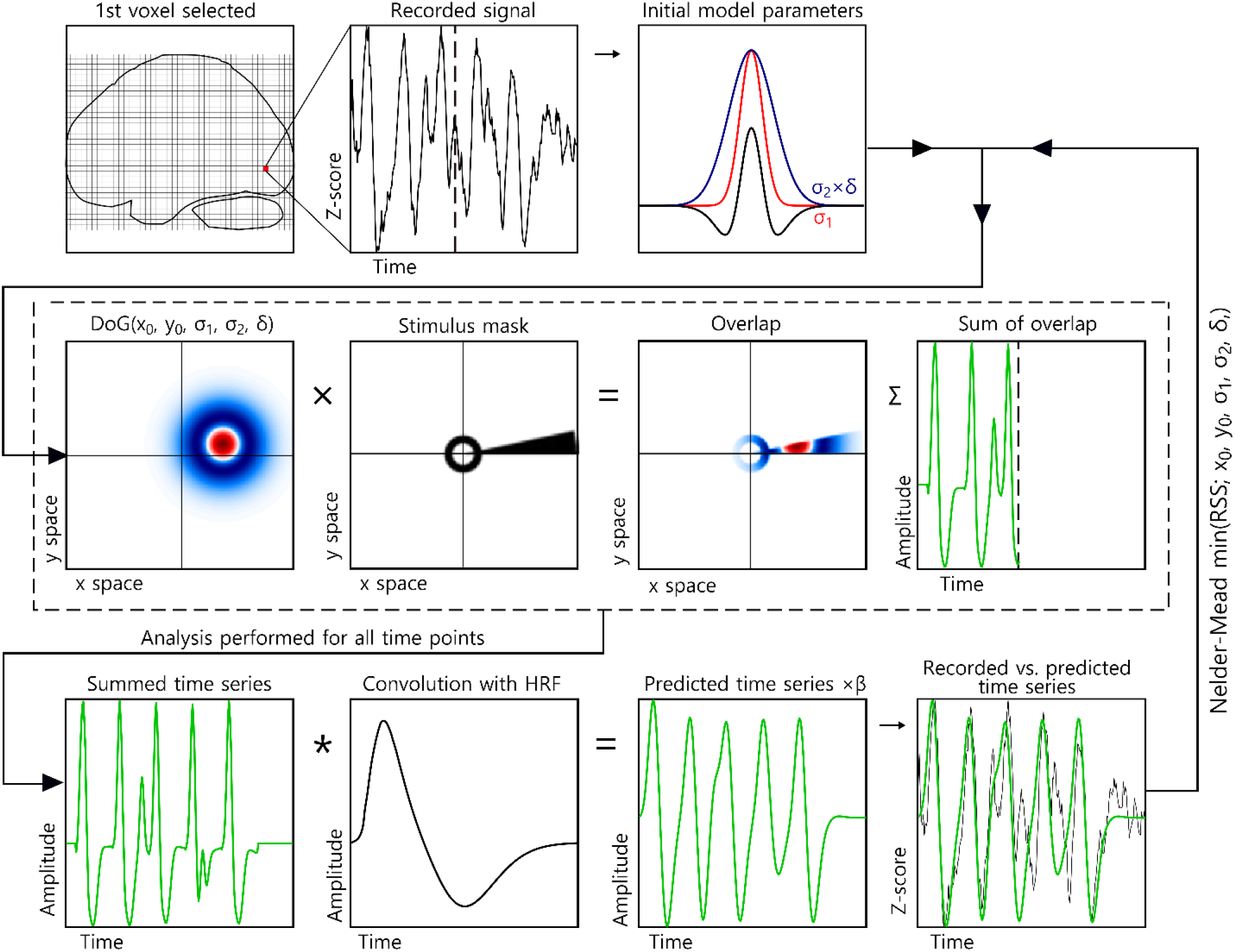
pRF mapping procedure. The time series of one voxel is selected and initial model parameters are obtained from a simple coarse fitting procedure. The overlap between these parameters is then multiplied with a binary mask of the stimulus sequence then summed for each TR. Convolution of the result and a haemodynamic response function is performed and scaled with β. This yields the final prediction for that particular combination of DoG model parameters, and this prediction is then compared with the recorded time series. Nelder-Mead minimization of RSS is then performed to find the best fitting DoG model parameters for each voxel.

This was then repeated for the entire time series for that voxel. This result was convolved with a haemodynamic response function to yield the prediction for the given DoG model parameters (x_0_, y_0_, σ_1_, σ_1_, δ) for a given voxels time-series. The error between the observed signal and the predicted time-series is then calculated using residual sum of squares (RSS). Using the Nelder-Mead algorithm, the RSS was then minimized by selecting new DoG model seed parameters, and the process was then repeated until a set of predetermined criteria were fulfilled. Finally, a scaling parameter, β, which accounts for the unknown unit of the BOLD signal, was found by solving a General Linear Model (GLM) that consisted of the measured time-series, the predicted time-series, an error term and the unknown β-value. We then evaluated model performance by comparing their coefficient of determination, R^2^, and Akaike’s Information Criterion (AIC) score.

### Model comparison

To evaluate model performance, we first investigated the amount of variance explained by each model using the coefficient of determination, R^2^, for both models and all voxels. We then calculated the mean R^2^ for each hemisphere and participant for voxels that had an R^2^ > 0.05. This result is plotted in Figure 3A. It appears the two models are equally capable of explaining the variance in the data. To see if this result was caused by a high number of poor fits, we performed the same analysis only including model predictions that had an R^2^ > 0.15 and R^2^ > 0.3. These results are plotted in Supplementary Figure S3. This figure also includes the adjusted coefficient of determination for the three R^2^ values. The same pattern from these two additional analyses is seen, i.e. the amount of variance explained is similar for the two models.

**Figure 3.**
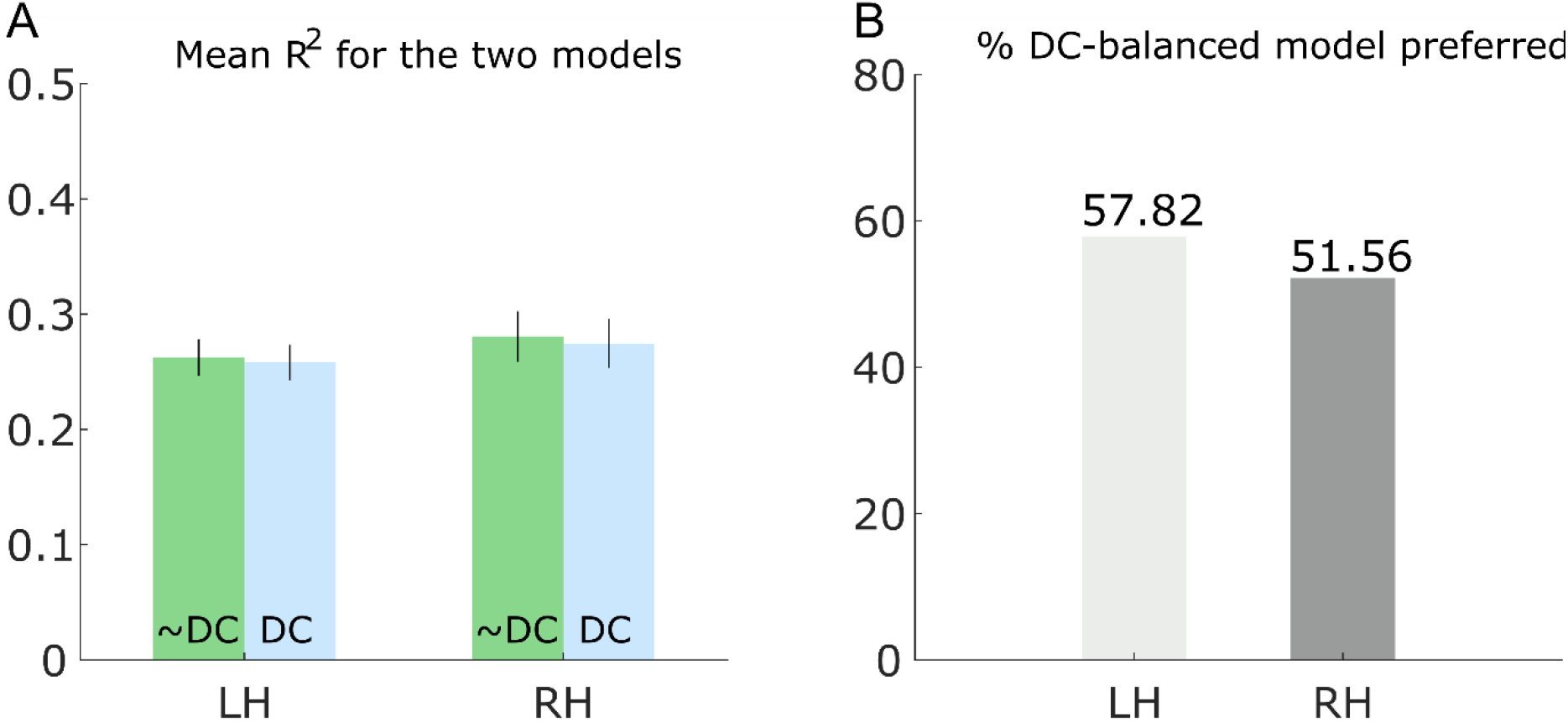
Comparison of R^2^ and AIC for the two models. **A:** Mean R^2^ for all participants’ left hemisphere (LH) and right hemisphere (RH) for the model without DC-balance restriction (~DC) and with DC-balance restriction (DC) when all voxels had had R^2^ > 0.05. The error bars indicate 95% confidence interval of the mean. **B:** Percentage of voxels in the two hemispheres that exhibited a lower AIC score for the DC-balanced DoG model.

To investigate whether the increased model complexity of the unrestricted model was justified, we further calculated Akaike’s Information Criterion (AIC)^13^. AIC penalizes models with a larger number of parameters and should here be a good indicator of whether the DC-balanced model with fewer parameters is preferable. AIC requires the compared models to be based on the same data. In this analysis, we therefore only included overlapping voxels from the two model predictions. For each participant, we calculated the AIC value for all voxels’ times-series (see methods for details). The model preferred is the one with the lowest AIC score. For each hemisphere, we then calculated the percentage of voxels where the DC-balanced model had the lowest AIC score. This result is plotted in Figure 3B, and here it can be seen that the DC-balanced DoG model had the overall best performance for both hemispheres. This suggests that the simpler DC-balanced DoG model is preferred over the unrestricted model when model complexity is taken into account.

### Original versus computed δ

Another method for examining if pRFs are DC-balanced is to examine the parameters of the unrestricted model. It is possible for this model to converge in a configuration that is highly similar or identical to the DC-balanced model without being forced to. In such a scenario with only very slight differences in parameter values, the extra parameter freedom is essentially non-utilised and the conclusion would be that even when unrestricted, the model is effectively DC-balanced. To examine if this was the case, we used equation 6 with σ_1_, σ_2_ and δ from the unrestricted model predictions to calculate the corresponding values that would result in a DC-balanced DoG. Comparing the original unrestricted model parameters to the computed parameters then gives an indication of how far the unrestricted model is from being DC-balanced. Figure 4 shows the three computed parameters plotted against the original for both the left and right hemisphere for all participants’ V1 voxels with an R^2^ > 0.05 and .1 < δ < .9.

**Figure 4.**
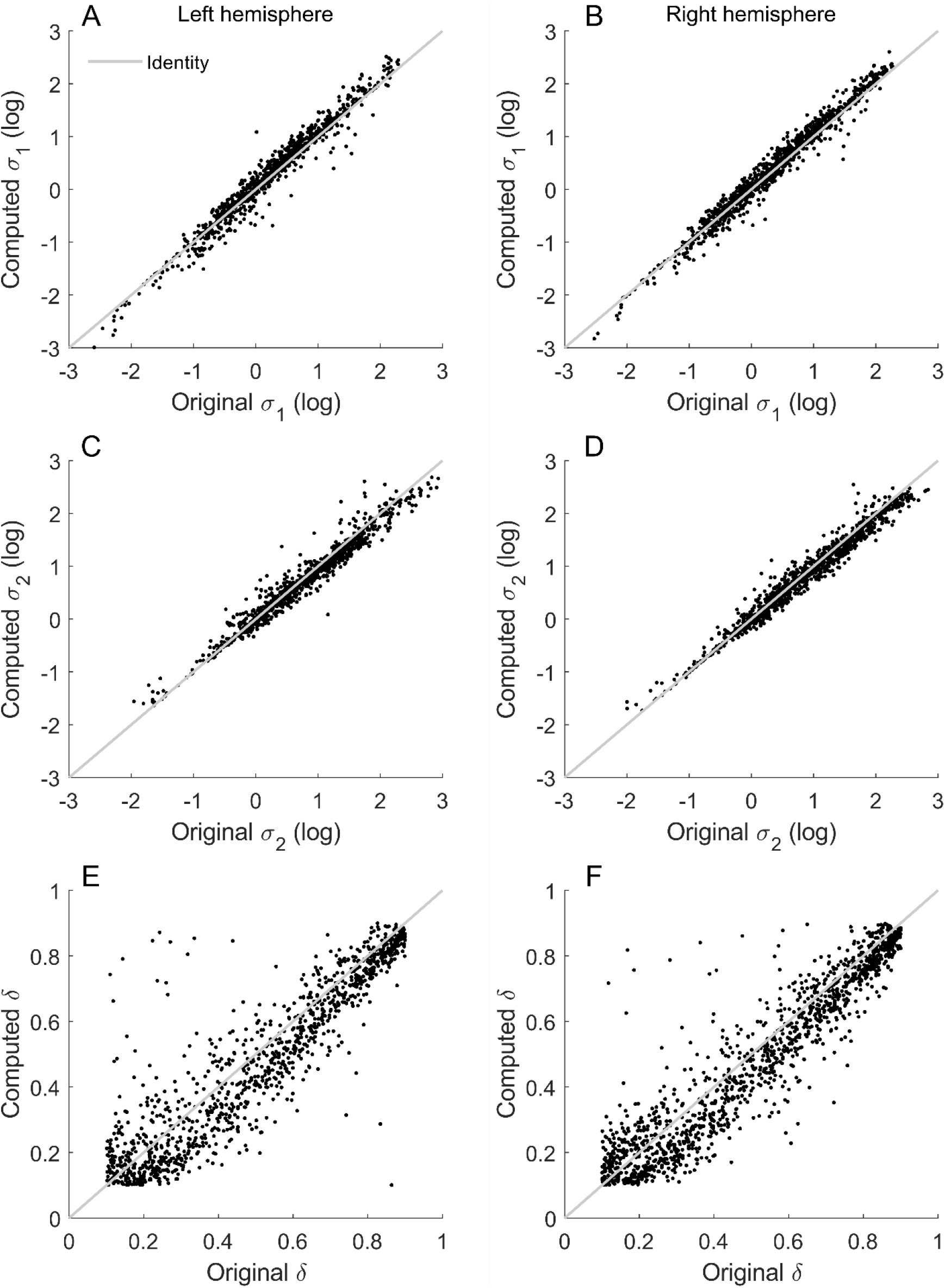
Comparison of fitted model parameters with computed model parameters according to equation 6. **A&B:** computed σ_1_ plotted against original σ_1_ for the left and right hemispheres. Light grey line shows the identity line. **C-F:** Same as A&B, but for σ_2_ and δ.

The σ_1_and σ_2_ has been log-transformed to accommodate for skewness of the data. The results are shown in Figure 4A-D, and it can be seen that the original and computed data lie close to the identity line. Figure 4E-F shows the results for δ. This data has not been log-transformed, so here deviations from the identity line are more noticeable, but a strong linear trend is still present. To compare the observed with the computed parameters, we used random coefficient models for each parameter separately. The results were highly significant for all parameters (Wald χ2(1) < 10000, p < 0.00001 in all cases). This suggests that the unrestricted model in practice resulted in model configurations that could be characterised as nearly or fully DC-balanced.

### Normalization

In most fMRI experiments, several preprocessing steps are applied before data analysis. Here, each voxel’s BOLD timeseries was detrended and Z-score normalized. The detrend operation is performed to remove what is known as signal drifts, which are low frequency signal oscillations that are thought to be associated with scanner drifts as well as cardiac and respiratory effects^14^. The detrend operation removes the best straight-line fit from the signal. Z-score normalisation also removes the mean of the signal but in addition also normalises the variance of each voxels timeseries with its standard deviation^15,16^. Z-score normalisation is applied to account for low-frequency variance not accounted for by the stimulus^17^. Because these normalisation steps may influence the two models’ performance, the model comparison was also performed on the same data, but without detrending and Z-score normalisation. These results are shown in Figure 5.

**Figure 5.**
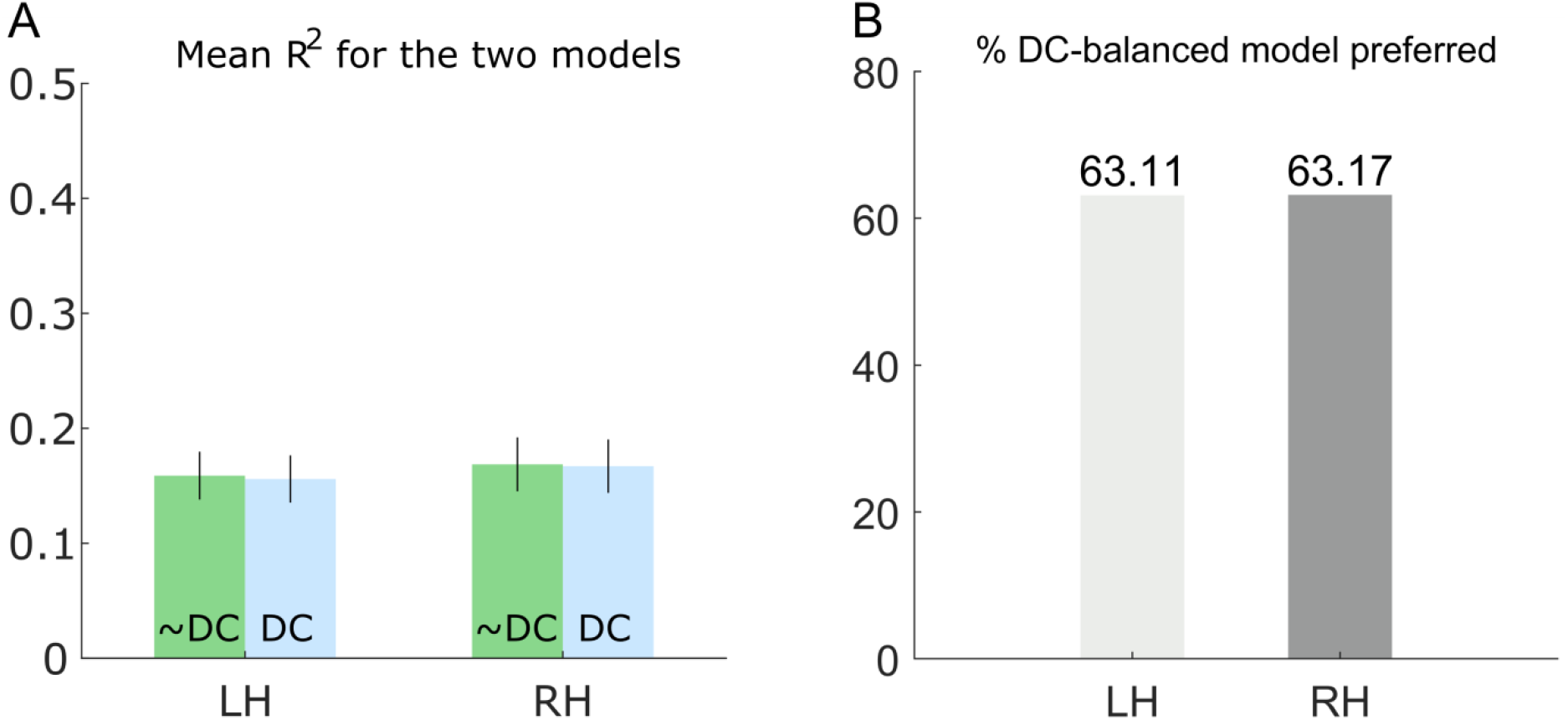
Comparison of R^2^ and AIC for the two models using non-normalised data. **A:** Mean R^2^ for all participants’ left hemisphere (LH) and right hemisphere (RH) for the unrestricted model (~DC) and with DC-balance restriction (DC) when all voxels had had R^2^ > 0.05 after being fitted to the non-normalized data. The error bars indicate 95% confidence interval of the mean. **B:** Percentage of voxels in the two hemispheres that exhibited a lower AIC score for the DC-balanced DoG model using the non-normalized data.

Figure 5A shows that the two models still perform similarly in terms of the amount of variance explained, but also that they explain less variance when compared to the normalised data (cf. Figure 3A). Figure 5B shows that the DC-balanced model is still favoured, but the DC-balanced model is now favoured in 63% of all voxels compared to ~55% for the normalised data (cf. Figure 3B). This suggests that both models benefit from normalisation in terms of the amount of variance they explain, but that the extra parameter of the unrestricted model is more beneficial when the data is normalised.

### DC-offset

In digital image analysis, the DC-offset of an image (or DC-component) is often thought to contain no interesting information^18^. A similar assumption is made in fMRI analysis when the BOLD signals’ DC-offset is removed by some form of mean subtraction (e.g. detrending or Z-score normalisation). However, as discussed in the introduction, the DC-offset is relevant for visual processing as this sets the scale for the variation in contrast. In the non-normalized dataset, the DC-offset of each voxels BOLD timeseries is captured by the constant term of the GLM and scaled with β. The scaling is applied on a voxel-by-voxel basis due to the unknown unit of the BOLD signal, but it might be speculated that a more accurate scaling could be extracted from the DC-offset. It is beyond the scope of this article to give a thorough examination of this question, but as a preliminary consideration, we investigated the relationship between the BOLD signal DC-offset and eccentricity in V1.

This was first done by back-projecting the DC-offset onto an inflated cortex for all voxels. This result is shown in Figure 6A for the left hemisphere and Figure 6B for the right hemisphere for a single participant, but the remaining participants showed similar results. These are shown in Supplementary Figure S4 and S5. It is immediately noticeable that there appears to be a negative gradient in the DC-offset as a function of eccentricity. To investigate this further, we looked at the DC-offset as a function of eccentricity for all participants and all voxels with an R^2^ > 0.05 within V1. Data were analysed in a random coefficient model with the DC-offset as the dependent variable, eccentricity (log-transformed) as a covariate and random intercepts and slopes varying with participant. Model fits were made using a restricted maximum likelihood function with the Broyden-Fletcher-Goldfarb-Shanno (BFGS) algorithm. The results were highly significant for both the left hemisphere (Wald χ2(1) = 82.76, p < 0.0001, intercept 95% CI: 309.56 to 339.22) and the right hemisphere (Wald χ2(1) = 175.19, p < 0.0001, intercept 95% CI: 289.39 to 313.03). We take these results to suggest there is an unexplored connection between the stimulus and the BOLD signal DC-offset in V1.

**Figure 6.**
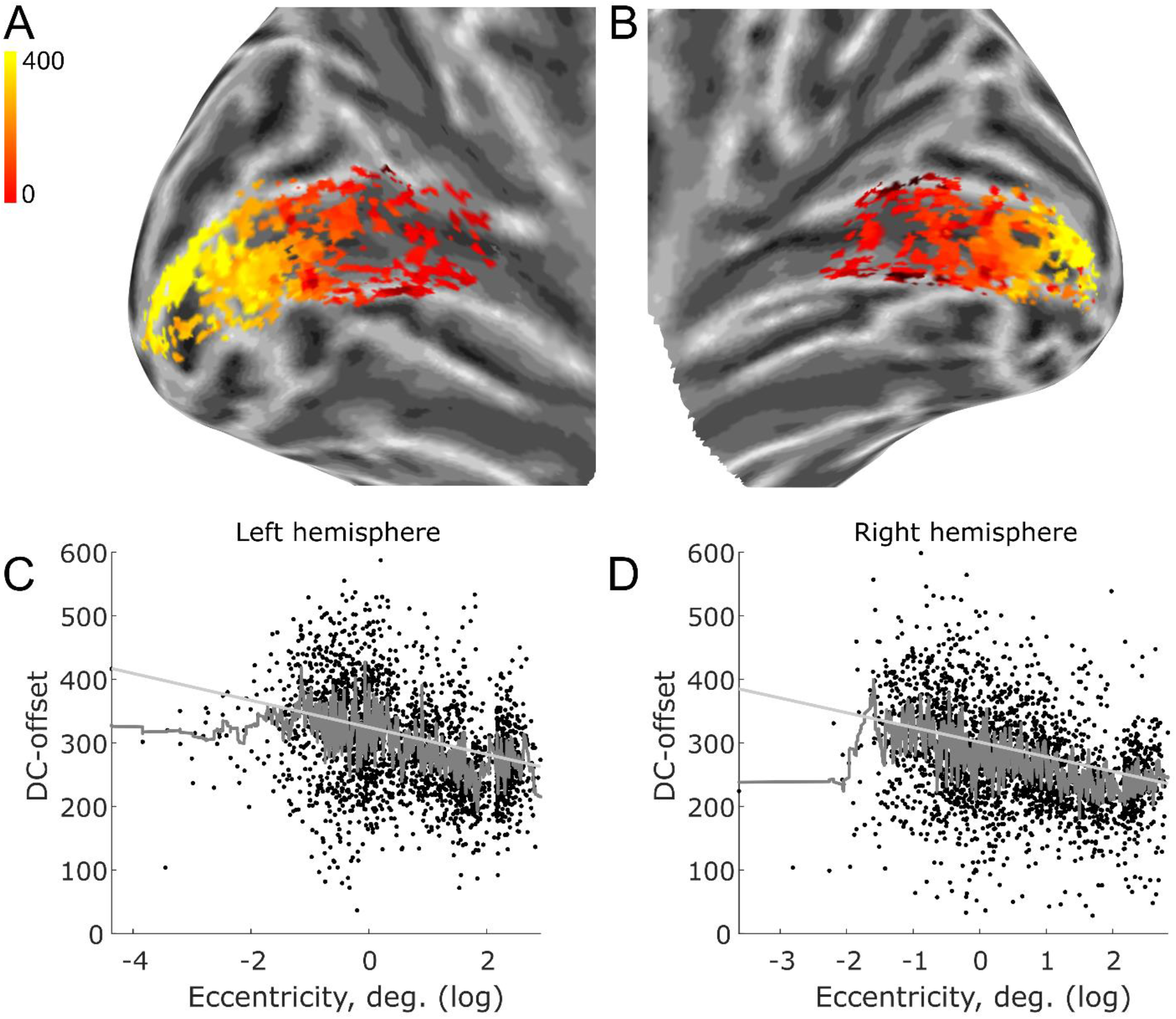
DC-offset as a function of eccentricity. **A&B:** Back-projection of the non-normalised DC-offset data for a single participants’ left and right hemisphere, respectively. **C&D:** Linear model (light grey) for DC-offset as a function of eccentricity (log scale) for left and right hemisphere, respectively. The dark grey line indicates the moving average with a window size of 100 voxels.

To demonstrate that the DC-offset is not caused by e.g. large positive deviations from a common offset between voxels, Figure 7 shows the maximum and minimum BOLD signal amplitude as a function of the DC-offset for voxels within V1 with an R^2^ > 0.05 for both hemispheres.

**Figure 7.**
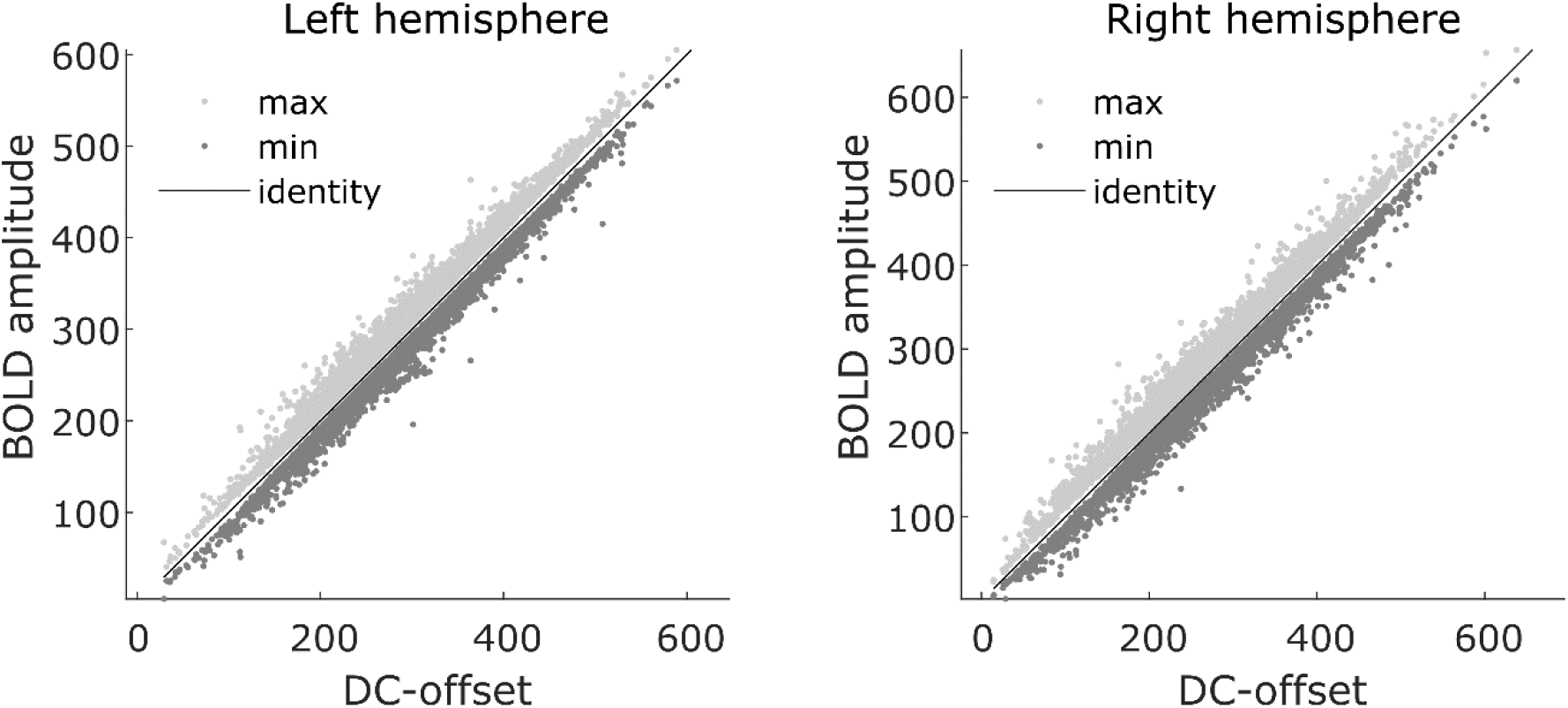
Maximum and minimum BOLD response as a function DC-offset. In both plots, the maximum BOLD amplitude is plotted as a function of DC-offset is plotted in light grey, whereas the minimum BOLD response as a function of DC-offset is plotted in dark grey. The identity line is shown in black.

For both hemispheres, it can be seen that the maximum BOLD response is consistently just above the identity line, whereas the minimum BOLD response is just below the identity line. This shows that a large DC-offset is accompanied by correspondingly large excitatory and inhibitory BOLD signal amplitude, meaning the BOLD signal varies around the DC-offset.

## Discussion

Prior research has suggest that the excitatory and inhibitory parts of the population receptive field tend towards DC-balance ^7^, and we believe our findings are in agreement with this. It should be noted, however, that this likely dependent on the specific stimulus protocol used. In this experiment, the stimulus protocol consisted of an expanding or contracting ring simultaneous with a clockwise or counter clockwise rotating wedge. This, however, may not be the optimal stimulus configuration to study DC-balance as it allows for situations where the ring and wedge enter the receptive field shortly after one another. In these cases, the post-stimulus undershoot of the haemodynamic response may not have had enough time to return to baseline before the next stimulation. This can result in a positive bias in the BOLD signal, albeit in our data it appears that these situations are not numerous enough to favour the unrestricted DoG model.

The DC-offset may also depend on the choice of stimulus. Since we presented our stimuli on a uniform grey background, epochs of no stimulation approximately correspond to the DC-offset (baseline). As illustrated in Figure 6, however, the DC-offset varies from voxel to voxel (in the non-normalised dataset). A negative relationship between DC-offset and eccentricity means voxels responding to foveal stimulation have a higher DC-offset compared to voxels responding to peripheral stimulation. Since the stimulus texture consisted of the same repeating 8Hz “rippling” texture up to 17.4 degrees visual angle (two frames of this texture are shown in Supplementary Figure S1 and S2), it could be speculated that this texture were the cause of the larger DC-offset in foveal voxels. This is because these receptive fields could be more sensitive to the spatial frequencies of the texture compared to the receptive fields of peripheral voxels. However, if this were the case, we would expect the DC-offset to relax to a common offset during the background epochs, but Figure 7 demonstrates that the BOLD signal remains close to the DC-offset throughout the timeseries. Alternatively, the DC-offset could be related to the cortical magnification factor (CMF). CMF is the distance on the cortical surface between two points representing visual field locations 1° apart^19^. CMF indicates that a higher number of neurons respond to one visual location compared to another, and CMF declines as a function of eccentricity from fovea to periphery^20^. The basic idea being that a foveal voxel containing more neurons results in overall higher mean BOLD response compared to a peripheral voxel. In addition to the CMF, as was also discussed in the introduction, it has been proposed that it is advantageous to sample stimulus contrast and luminance separately. In line with this reasoning and if the receptive fields are indeed DC-balanced, it may be proposed that the variance modelled by the DC-balanced DoG is the contrast of the stimulus, while the DC-offset may be closer related to stimulus luminance. Stimulus luminance has long been associated with maintained neural activity^21,22^. It therefore seems natural to suggest that the maintained activity may also be related to the DC-offset of the fMRI BOLD response, i.e. a higher maintained firing rate yields a higher mean BOLD response. Note that this effect does not exclude the CMF from also playing a role in the BOLD signal DC-offset. In the work that originally proposed the DoG as a receptive field model^23^, the full model also treated the maintained and transient components separately. Here, albeit without explicitly stating so, it is seen that the transient component models stimulus contrast and the maintained component models stimulus luminance. Future research could investigate this in further detail by manipulating both stimulus contrast and luminance, and see if it is possible to model them separately.

In our model comparison, we did not include the single Gaussian pRF model since this model has already been compared to the unrestricted DoG^5^. Furthermore, the single Gaussian model relies heavily on a haemodynamic response function (HRF) with a large post stimulus undershoot (PSU), as it itself is not capable of modelling the parts of the BOLD signal below the DC-offset. In this experiment, we have also used a custom HRF that displays a large PSU (see Figure 2). This HRF was measured empirically in the context of a pRF mapping experiment^24^. Since we used the same HRF for both the unrestricted and the DC-balanced model, both models can take advantage of the larger PSU. It might, however, be the case that this HRF aids the unrestricted model when it is positively biased, since this HRF can supply a source of inhibition that it would otherwise not have access to. In general, it may be problematic that the relationship between stimulus and BOLD signal is modelled first with the pRF model and subsequently with the HRF, as this makes the relationship between the pRF model and the stimulus difficult to interpret. The scaling with β applied in the general linear model further complicates this relationship, since the sensitivity of a given receptive field is now also dependent on β. In future studies, it would therefore be beneficial to investigate how the choice of HRF affects model predictions. Furthermore, it may also be questioned whether pRF modelling in the framework of a general linear model is the most appropriate method. This, however, requires a better understanding of the scaling parameter, β, which is notoriously difficult to model since it is likely to be dependent on the adaptive mechanisms of the visual system. The DC-offset of the BOLD signal could perhaps also aid in this endeavour.

## Conclusions

We have determined the conditions for a DoG to be DC-balanced and subsequently examined whether pRFs of V1 can be said to be DC-balanced. By considering model complexity with AIC, it was shown that the DC-balanced model was preferred. We next showed that the unrestricted model is close to being DC-balanced by comparing the model parameters of the unrestricted to model parameters that were computed by enforcing DC-balance. To ensure our findings were not due to normalisation, we also conducted the model comparison on non-normalised data and found that the DC-balanced model was still preferred over the unrestricted model. These results indicate that V1 neurons are at least frequently organised in the exact constellation that allows them to function as bandpass-filters, which can be considered the constellation that allows for the complete separation of processing contrast and luminance information. Finally, we showed results suggesting that the DC-offset of the V1 BOLD signal may contain information relevant to the neural processing of visual information. We speculate that this visual information may correspond to the stimulus luminance. If this is the case, it may be possible to separate luminance and contrast processing in fMRI images of the visual system, which could lead to a new understanding of the haemodynamic response.

## Methods

### Participants

This study was approved by the local ethics committee, De Videnskabsetiske Komitéer for Region Midtjylland, and conducted according to the principles of the Declaration of Helsinki. 24 healthy, right-handed adults with normal or corrected-to-normal visual acuity aged between 18 and 40 (14 females) gave informed consent to participate in the study. 2 participants were excluded because we failed to generate a meaningful pRF map for either one or both hemispheres.

### MRI and stimuli

The stimuli were similar to those used in conventional retinotopic mapping^25^ and consisted of expanding/contracting rings and clockwise/counterclockwise rotating wedges with a high-contrast ripple-like texture and mean luminance of 87 cd/m^2^. The stimuli subtended a maximum of 17.4° visual angle. Example frames of this stimulus texture are provided in Supplementary Figures S1 and S2. The stimulus texture was updated at 8hz.

### Procedure

Participants were placed in the bore of a Siemens Magnetom Skyra 3T MRI scanner. Stimuli were projected onto a screen in the back of the scanner and viewed by the participant through a mirror placed on top of the head coil. Participants were instructed to maintain fixation on a small black dot in the centre of the screen and at pseudo-random intervals, the dot would change colour to red. To control eye-fixation and overt attention, participants were instructed to press a button when this occurred.

Stimulus presentation and recording of behavioural responses was implemented using the Psychophysics Toolbox 3^26,27^ running under MATLAB 2019b. Population Receptive Field mapping was done using the SamSrf toolbox and custom scripts. fMRI images were acquired using a 32-channel head coil with the top removed (for better participant view) resulting in 20 remaining channels.

176 whole-brain volumes of high-resolution structural images were acquired using a T1-weighted coronal-oriented MP2RAGE sequence (TR: 5000ms, TE: 2.98ms; field of view: 240 x 256 x 176mm, voxel size: 1mm^3^ isotropic) using a 32-channel head coil.

The pRF mapping protocol acquired 15 continuous axial slices (slice-thickness: 2mm) for each volume using a T2*-sensitive gradient echoplanar Imaging (EPI) sequence (TR: 1000 ms; TE: 30 ms; flip angle: 60°; field of view: 192 x 192 x 30 mm; voxel size: 2×2×2 mm, GRAPPA with PAT-factor 2). 6 runs of 225 volumes were acquired for each participant (3 clockwise and 3 counter-clockwise).

### MRI analysis

Image preprocessing was carried out using Statistical Parametric Mapping (SPM12; Wellcome Department of Imaging Neuroscience, London, UK) running under MATLAB R2019b. For each participant, volumes from the pRF mapping were realigned to the first image of the run to correct for head motion, unwarped using a presubstracted phase and magnitude B_0_ field map to correct for geometric distortions, and slice-time corrected for acquisition delays. Next, the first image of the first run was coregistered to the high-resolution structural and subsequently these coregistration parameters were applied to all other images from the remaining runs. The 3 clock-wise runs and 3 counter clock-wise runs were averaged separately. Thus, there were 2 averaged runs for each participant and these were concatenated so that the complete timeseries consisted of 450 images. Prior to the first pRF analysis, each voxels’ timeseries was detrended and Z-score normalised. These steps were omitted for the second pRF analysis (non-normalised data). No spatial or temporal smoothing was performed. The structural images were used for cortical reconstruction and volumetric segmentation within the Freesurfer image analysis suite^28,29^ to create an inflated 3D image of the grey/white-matter boundary for each participant. Based on this reconstruction, a prediction of the anatomical extent of V1 was created using a built-in Freesurfer algorithm^11^. This V1 delineation was used as a mask for constraining which voxels to include in the following pRF analysis.

### pRF analysis

A full account of the pRF analysis procedure has been given by Dumoulin and Wandell^5,9^. To aid us in this analysis, we used the pRF mapping toolbox ‘SamSrf’ developed in the lab of Sam Schwarzkopf. The toolbox can be found at https://sampendu.net/. In brief, we modelled the measured fMRI signal as

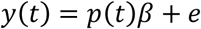

where *p*(*t*) is the predicted BOLD signal in each voxel at time *t*, *β* is a scaling coefficient that accounts for the unknown unit of the BOLD signal and *e* is the residual error. The predicted response *p*(*t*) was calculated by modelling the receptive field of the neuronal population within each voxel using a two-dimensional Difference of Gaussians of the form

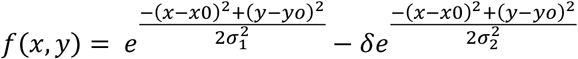

where σ_1_ and σ_2_ is the standard deviations of the two Gaussians and δ is a parameter that modulates the amplitude of the second negative Gaussian. This model was multiplied with a binary matrix representation of the stimulus intensity at (x,y) for all time points in the acquired volumes. The result was summed for each time point and the resulting timeseries was then convolved with the SAM Surfer toolbox default hemodynamic model, and finally scaled with β. We used Matlab’s implementation of the Nelder-Mead method (fminsearch) to obtain the pRF model parameters that best fit the data by minimizing the sum of squared error between the observed and predicted response.

### Model comparison

**The coefficient of determination**, R^2^, was calculated according to

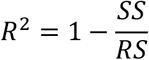

where SS is the sum of squares and RS is the residual sum of squares and defined as

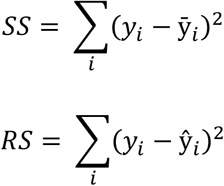

where y_i_ is the i^th^ time-series of the observed data, 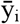 is the mean of the i^th^ observed time-series, and 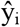 is the i^th^ times-series of the fitted data.

**Akaike’s Information Criterion** was calculated as follows:

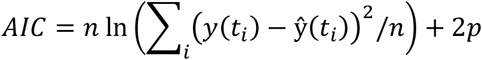

Here, *n* is the number of observations, *p* is the number of parameters, *y* is the measured BOLD signal, 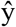 is the fitted values for the two models, and *t*_*i*_ is the i^th^ element of the time-series.

**The adjusted coefficient of determination**, 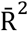 was based on R^2^, but adjusted so that

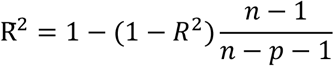

where *p* is the number of parameters and *n* is the sample size. These results are reported in Supplementary Figure S3.

