## Supplementary Information for "Population receptive fields of human primary visual cortex organised as DC-balanced bandpass filters"

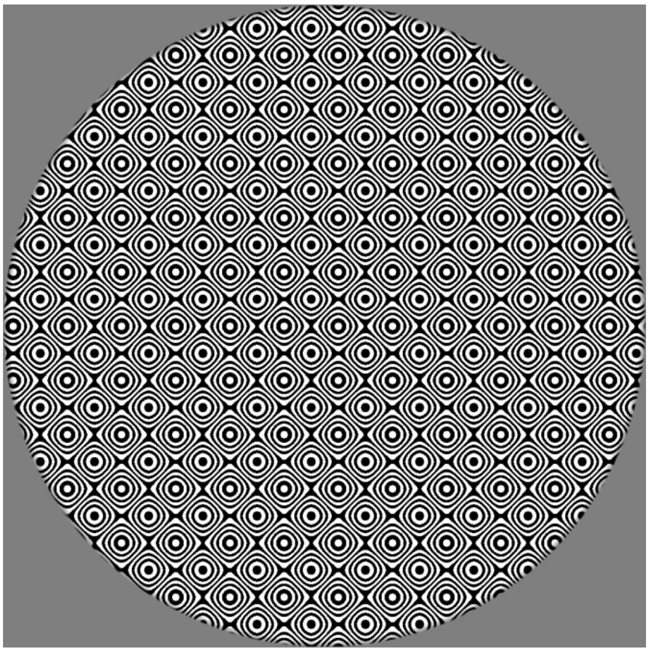

**Supplementary Figure S1. Example pRF stimulus texture frame.** One frame of the stimulus texture used in the pRF experiment shown at 30% of its original size.

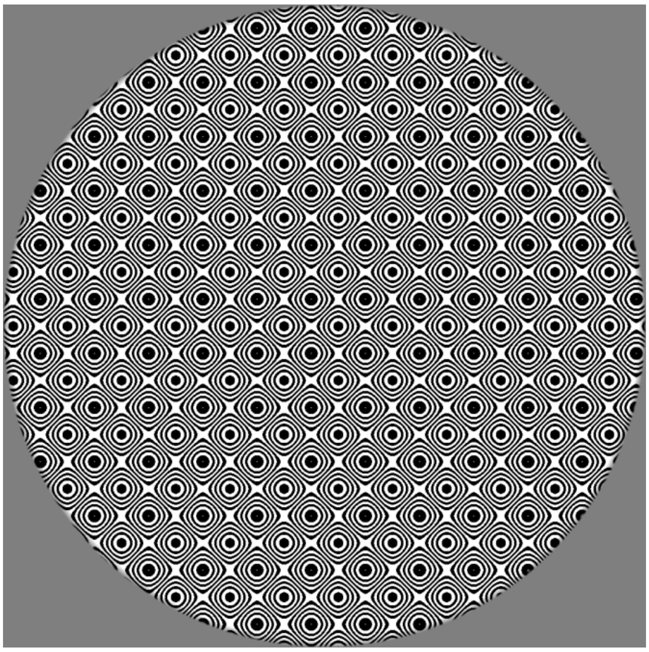

**Supplementary Figure S2. Example pRF stimulus texture frame.** Another texture frame from the same pRF stimulus experiment shown at 30% of its original size.

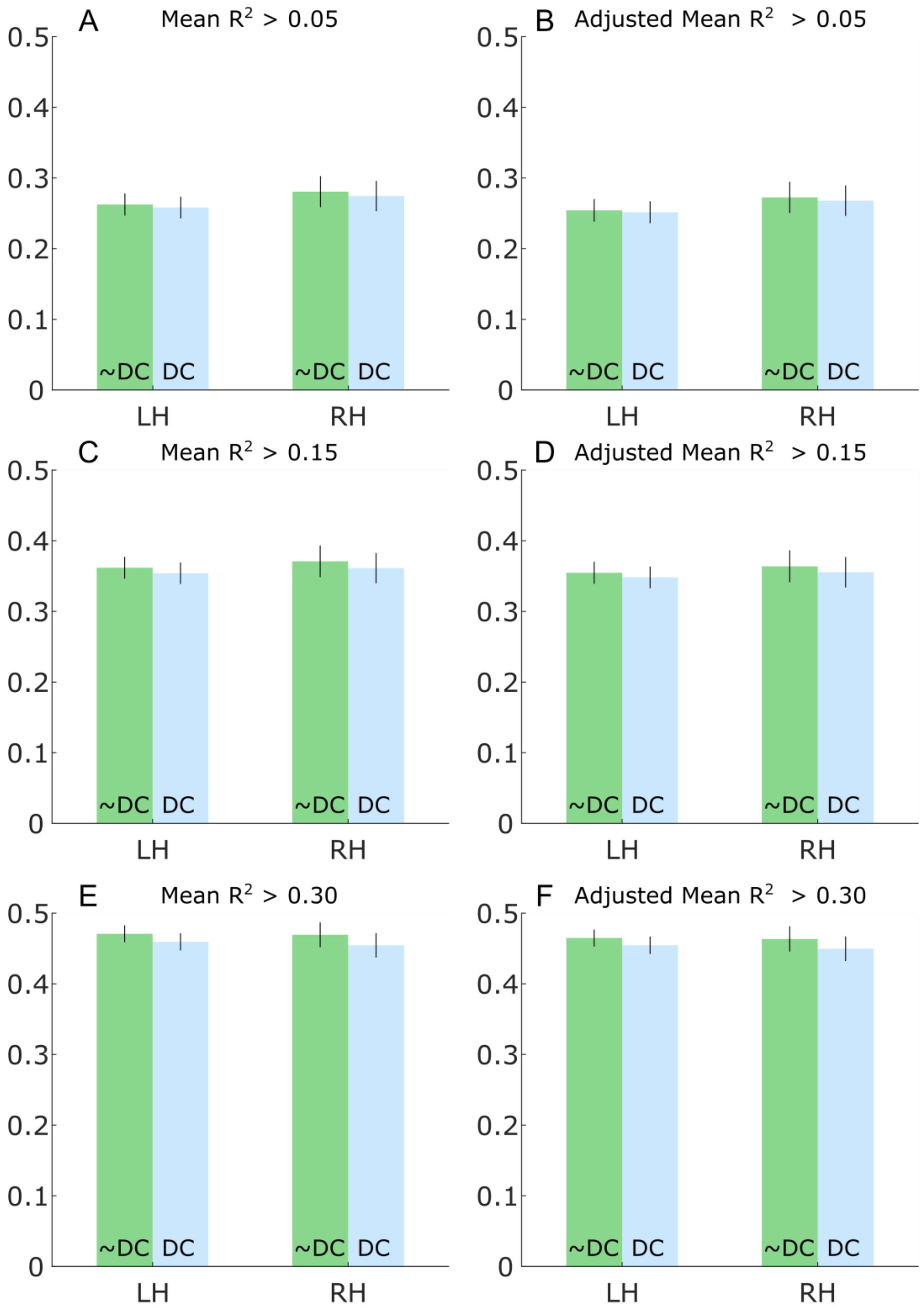


**Supplementary Figure S3. Comparison of $R^2$ and adjusted $R^2$ for the two models with varying $R^2$. A:** Mean $R^2$ for all subjects left hemisphere (LH) and right hemisphere (RH) for the model without DC-balance restriction (~DC) and with DC-balance restriction (DC) when all voxels had $R^2 > 0.05$. The error bars indicate 95% confidence interval of the mean. **B:** Same as A, but for adjusted $R^2$. **C-D:** Similar to A and B, but for $R^2$ and adjusted $R^2 > 0.15$ and 0.3, respectively.

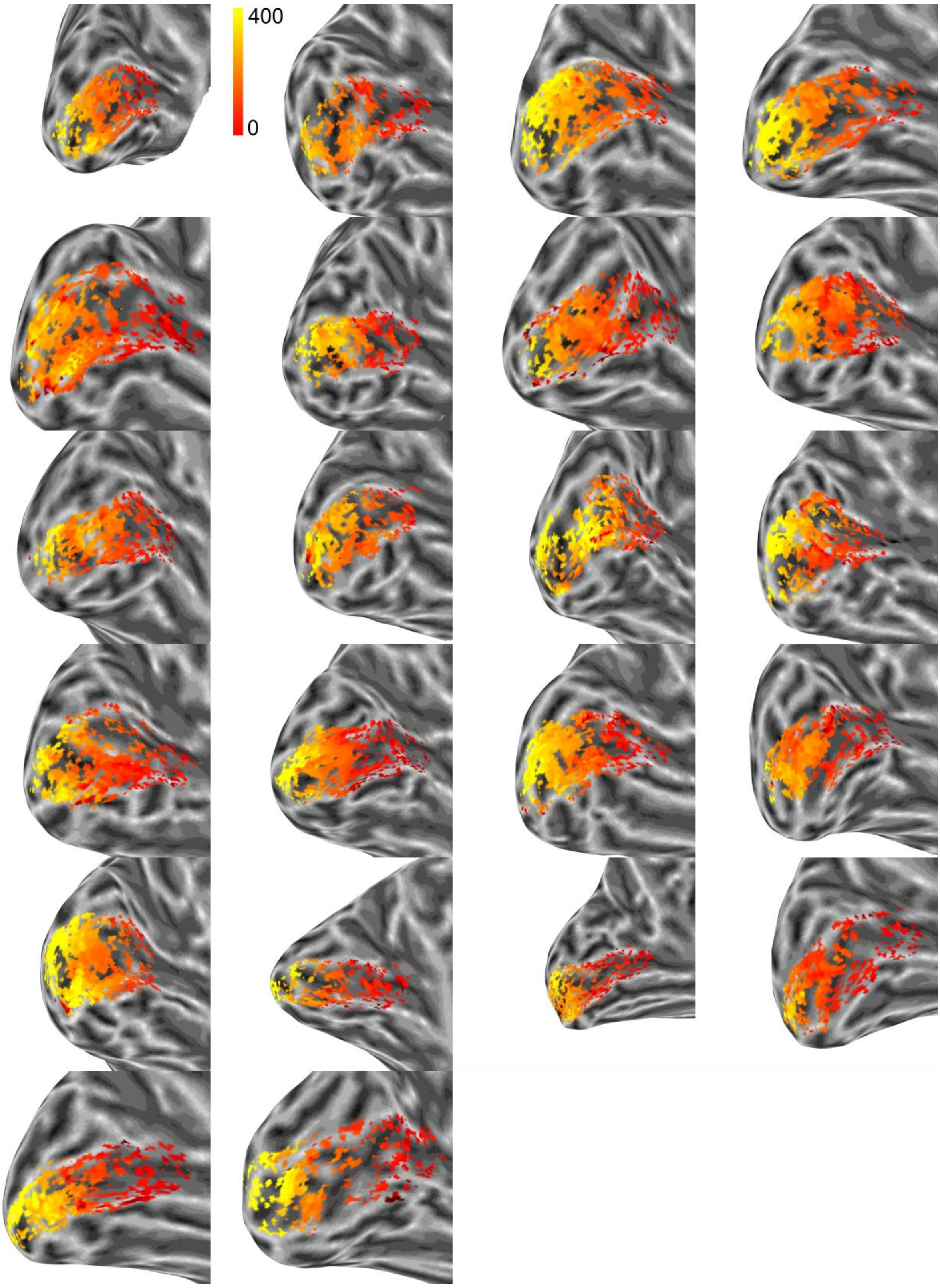


**Supplementary Figure S4. DC-offset as a function of eccentricity**. Back-projection of the non-normalised DC-offset data for all 22 participants' left hemisphere.

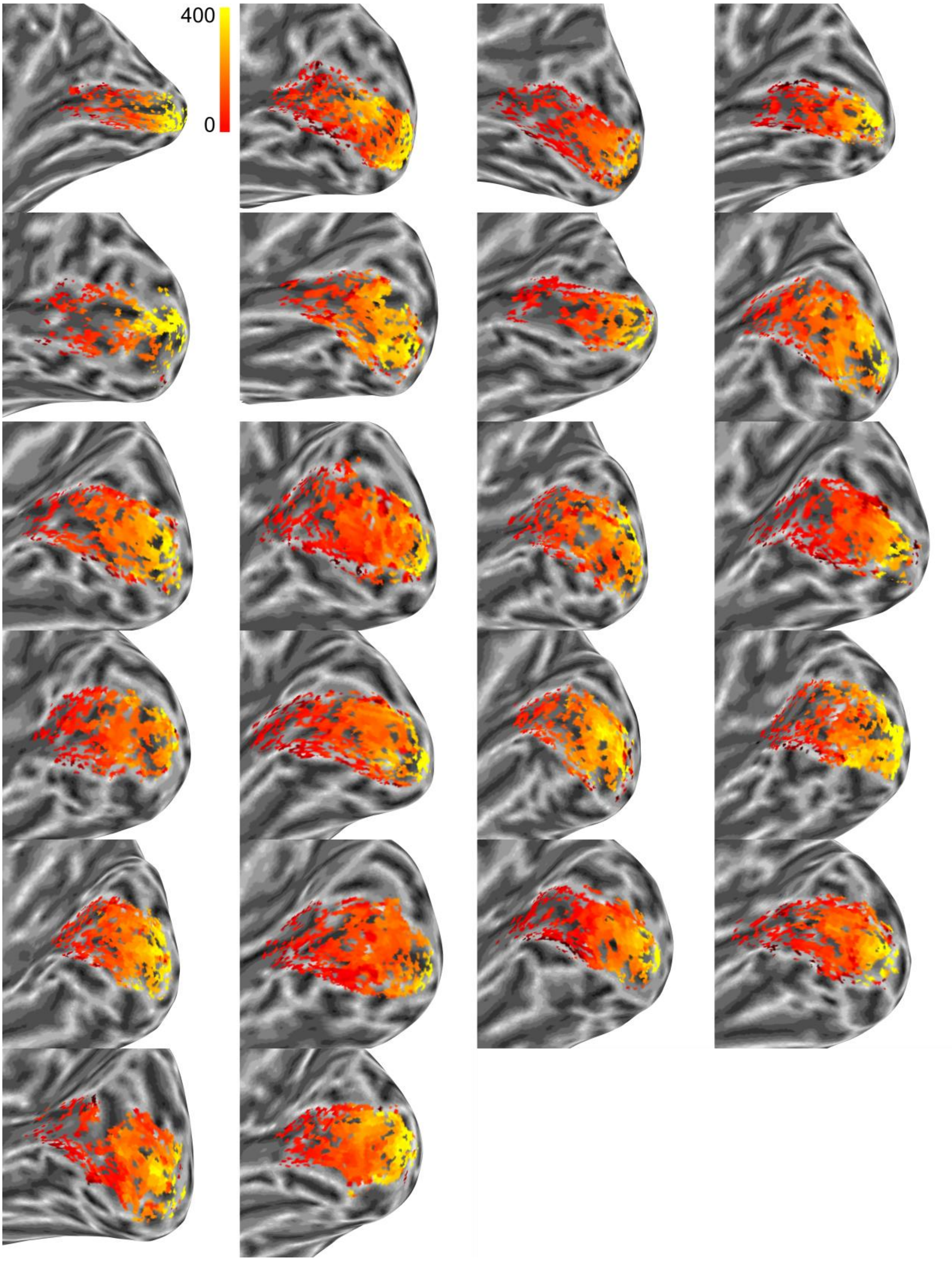


**Supplementary Figure S5. DC-offset as a function of eccentricity.** Back-projection of the non-normalised DC-offset data for all 22 participants' right hemisphere.
